# Predicting cell type-specific extracellular vesicle biology using an organism-wide single cell transcriptomic atlas – insights from the *Tabula Muris*

**DOI:** 10.1101/2024.02.19.580983

**Authors:** Thomas J. LaRocca, Daniel S. Lark

## Abstract

Extracellular vesicles (EVs) like exosomes are functional nanoparticles trafficked between cells and found in every biofluid. An incomplete understanding of which cells, from which tissues, are trafficking EVs *in vivo* has limited our ability to use EVs as biomarkers and therapeutics. However, recent discoveries have linked EV secretion to expression of genes and proteins responsible for EV biogenesis and found as cargo, which suggests that emerging “cell atlas” datasets could be used to begin understanding EV biology at the level of the organism and possibly in rare cell populations. To explore this possibility, here we analyzed 67 genes that are directly implicated in EV biogenesis and secretion, or carried as cargo, in ∼44,000 cells obtained from 117 cell populations of the *Tabula Muris*. We found that the most abundant proteins found as EV cargo (tetraspanins and syndecans) were also the most abundant EV genes expressed across all cell populations, but the expression of these genes varied greatly among cell populations. Expression variance analysis also identified dynamic and constitutively expressed genes with implications for EV secretion. Finally, we used EV gene co-expression analysis to define cell population-specific transcriptional networks. Our analysis is the first, to our knowledge, to predict tissue- and cell type-specific EV biology at the level of the organism and in rare cell populations. As such, we expect this resource to be the first of many valuable tools for predicting the endogenous impact of specific cell populations on EV function in health and disease.

## BACKGROUND/RATIONALE

Extracellular vesicles (EVs) like exosomes are cell-derived particles that support robust and dynamic intercellular and interorgan communication. EVs have been implicated in a variety of human diseases and health-promoting adaptations (e.g., exercise), but mechanistically linking a disease phenotype to EVs from specific cell types or tissues remains a grand challenge in the field of EV biology [1]. This knowledge gap exists due to the complexity of EV biogenesis, trafficking and composition. For example, it is widely accepted that most, if not all, cell types can secrete EVs, but this oversimplification of EV biology does not account for heterogeneity in the amount of EVs secreted by different tissues and cell populations. Our group and others have shown that EV secretion differs greatly (∼100x) across different cell types and tissues [2, 3], and for this reason, heterogeneity of EV secretion is clearly a relevant factor to understand EV biology at the level of the organism. Furthermore, EV composition is dependent at least in part upon cellular expression of EV cargo genes and proteins, but the same cargo (e.g., tetraspanins) can be found in EVs secreted from many different cell types. For example, “hallmarks” of EVs (i.e., CD9, CD63 and CD81) label different EV populations in the blood [4] and are differentially secreted by different cell populations [3], highlighting the need for a more comprehensive understanding of cell population expression of these genes and proteins. Finally, the vast majority of what we understand of EVs comes from cell populations that are naturally abundant and/or easy to culture, while EVs from rare cell populations are almost entirely undefined. The ubiquitous origins of EVs combined with their underlying but poorly defined heterogeneity are barriers to our understanding of EV biology in health and disease.

Functional associations exist between the secretion of specific EV cargo and the originating cell’s expression of genes and proteins involved in EV biogenesis and secretion. For example, our group has recently shown that expression of EV genes and proteins predicts tissue-specific EV secretion *ex vivo* [2]. Others have demonstrated that expression of EV proteins in a variety of immortalized cell lines predicts their incorporation into secreted EVs [3]. Applying these principles, here we predict which cell types may have the most robust capacity to secrete EVs and specific EV cargo by creating a transcriptomic “map” of EV-associated gene expression using the *Tabula Muris* [5] – a public single-cell RNA-Seq dataset consisting of 44,000 cells across 117 cell populations from 20 mouse tissues. While there are thousands of proteins and RNAs that may be trafficked by EVs, in this project, we carefully selected 67 genes involved in EV biogenesis, cargo sorting, secretion and cargo based on recent seminal reviews [6, 7] and guidelines from leading societies in the field of EV biology [8]. Our intent is to generate a resource built upon foundational EV genes that can be expanded upon in future studies.

## RESULTS

### Identification of EV genes for analysis

In this report we focused on genes that code the machinery required for EV biogenesis/secretion, protein sorting machinery and the most abundant protein cargo (**Figure 1**). The biogenesis and secretion of both exosomes and ectosomes (microvesicles) can be initiated through the recruitment of endosome-sorting complex required for transport (ESCRT) machinery [7, 9]. ESCRT-dependent processes begin with the sequential recruitment of ESCRT-0, I, II and III to an inward (exosome) or outward (microvesicle) budding region of cell plasma membrane. ESCRT-0 (Arrdc1, Gag, Hgs, Pdcd6, Stam and Stam2) and ESCRT-I (Mvb12[a and b], Tsg101, Ubap1, Vps28 and Vps37[a, b, c and d]) complexes possess ubiquitin-binding domains that aggregate ubiquitylated protein cargo via condensation, select a region of the plasma membrane that will become the resulting endosome and initiate membrane curvature [10–12].

**Figure 1:**
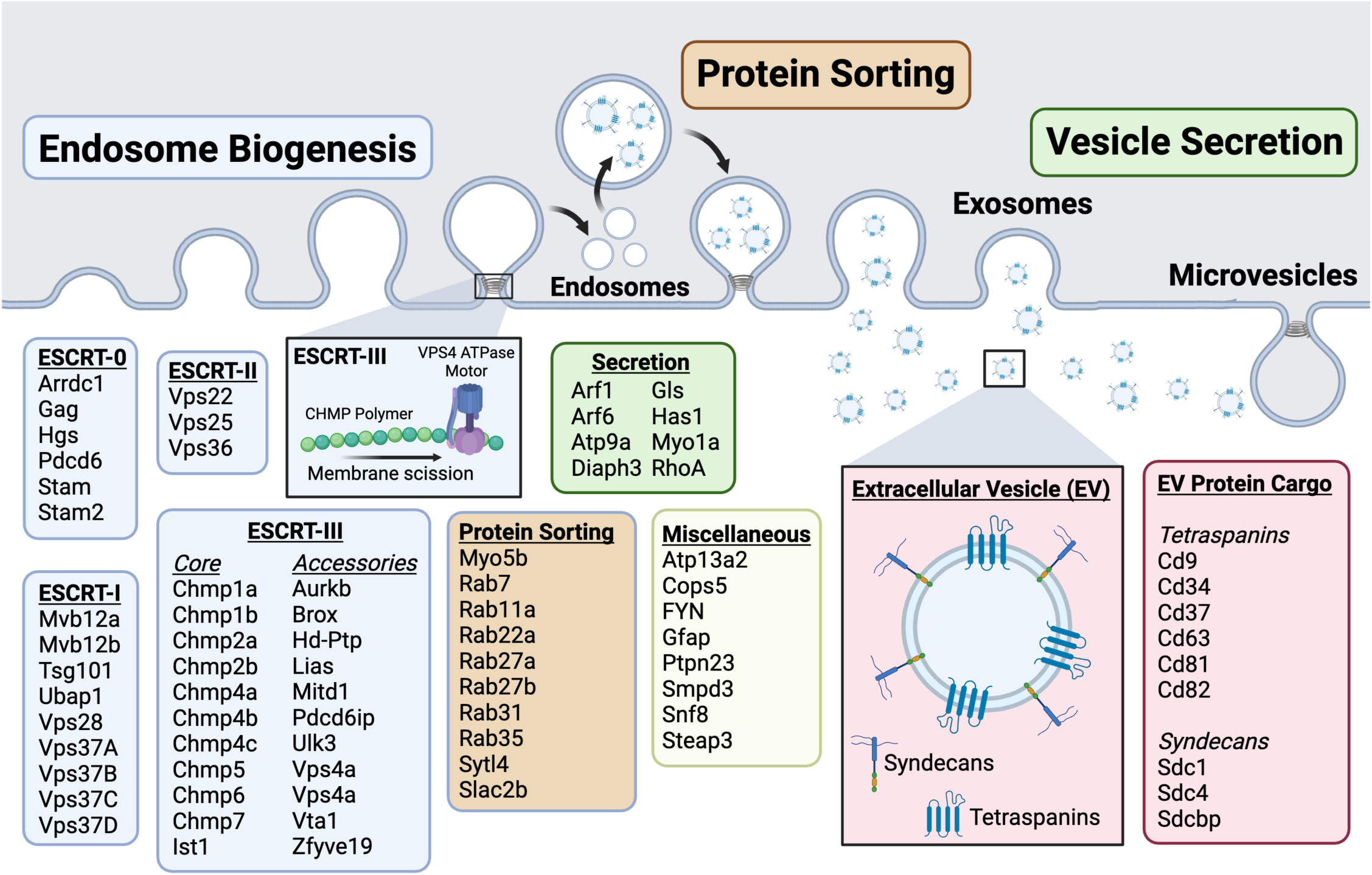
Overview of genes selected based on links to endosome biogenesis, endosome protein sorting, secretion of exosomes and microvesicles and common extracellular vesicle (EV) protein cargo. Made using Biorender.

ESCRT-II (Vps25, Vps22 and Vps36) sorts proteins within the membrane region forming the endosome [13, 14] and assembles ESCRT-III [15]. ESCRT-III is a filament (polymer) of charged multivesicular body proteins (Chmps) including Chmp1a, Chmp1b, Chmp2a, Chmp2b, Chmp4a, Chmp4b and Chmp4c, which wrap around the inward curving region of the membrane [16, 17]. Other Chmp family members with putative roles in ESCRT-dependent processing include Chmp3, Chmp5, Chmp6, Chmp7, Chmp8 and Ist1 [18]. ESCRT-III filaments are “pulled” and cleaved by a group of ATPase-driven molecular motors (Vps4a and Vps4b) with its co-factor (Vta1) and other related modulatory proteins (Brox, Lips, Anchr, Aurkb, Ulk3, Mitd1) to facilitate membrane closure and scission [19, 20]. While the ESCRT machinery is the most heavily studied mechanism of EV biogenesis, it is important to note that EVs can be formed independent of this specific process [21–24] and that the ESCRT machinery is also used for other cellular processes (e.g., cytokinesis)[6].

After endosome formation, vesicles mature within multivesicular bodies (MVBs) where protein cargo can be sorted by Rho GTPases and their co-factors (Myo5b, Rab7, Rab11a, Rab22a, Rab27a, Rab27b, Rab31, Rab35, Slac2b and Slp4) [25–28]. EV cargo can also be modified by Cops5 [29]. Other interesting EV-associated genes include: Atp13a2, FYN, Gfap, Ptpn23, Smpd3, Snf8 and Steap3. The cargo found within secreted EVs includes many of the proteins involved in biogenesis and secretion, along with a variety of other membrane embedded, intraluminal and extravesicular (“corona”) proteins. Inconsistent methodologies and technical challenges have led to disagreement regarding which proteins are physically associated with EVs *in vivo*. Nonetheless, genetic and proteomic studies of EVs in circulating blood plasma have found that tetraspanins (Cd9, Cd63, Cd81 and Cd82), syndecans (Sdc1, Sdc4) and syntenin-1 (aka Sdc binding protein (Sdcbp)) are amongst the most abundant EV proteins [30, 31]. The tetraspanins Cd34 and Cd37 have also been linked to EVs. Here, we studied the cell population-specific expression of the genes listed above to generate new insight and offer new hypotheses into cell type-specific EV biology applicable to a range of biologically important questions.

### The most abundant EV-associated genes encode the most abundant circulating EV proteins

We first identified abundantly expressed EV genes by plotting mean transcript abundance (FPKM) against expression frequency (% of cells > 50 FPKM) for every cell regardless of cell population. Seven EV genes (out of 67 analyzed) had abundant expression across the entire dataset (**Figure 2A**). These genes encode tetraspanins (Cd81, Cd9 and Cd63), syntenin (Sdcbp), syndecan 4 (Sdc4), Rho kinase A (RhoA) and adenosine diphosphate ribosylation factor 1 (Arf1). Five of these genes (Cd9, Cd63, Cd81, syntenin and Sdc4) encode some of the most abundant EV proteins secreted by cells in culture [30] and found in circulating blood plasma [31]. Moreover, a landmark study established that Sdcbp expression directly promotes the secretion of tetraspanin-expressing EVs [21]. Our analysis here is the first evidence that the natural abundance of these five EV proteins in the circulation could be due to high gene expression across many cell populations.

**Figure 2:**
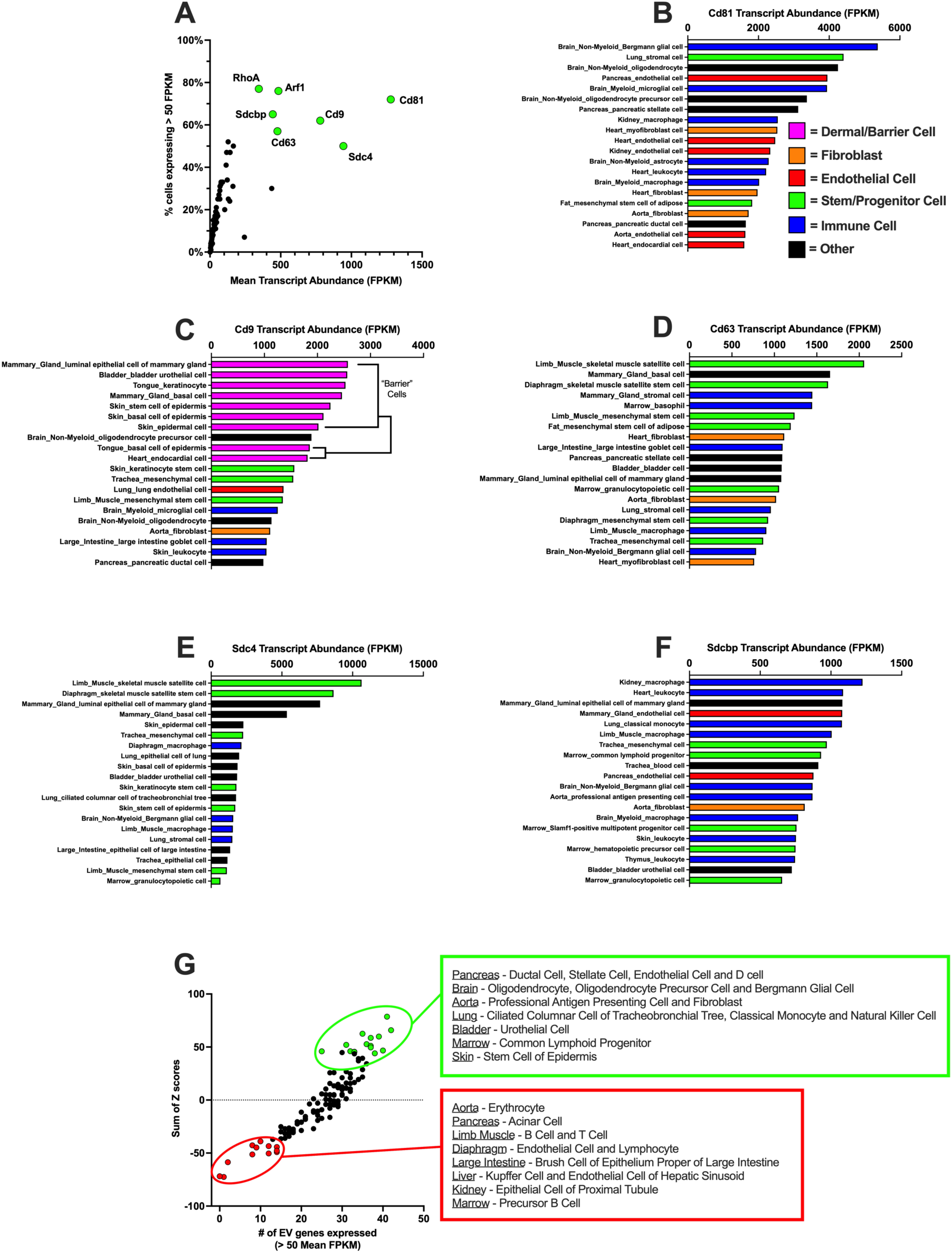
The most abundant EV-associated genes encode the most abundant circulating EV proteins but are enriched in different cell populations. A) Mean mRNA expression (FPKM) was plotted against the % of cells expressing at least 50 FPKM. Seven abundant and frequently expressed genes were identified. Five (Cd81, Cd63, Cd9, Sdc4 and Sdcbp) encode abundant EV protein cargo and were studied further. The top 20 cell populations expressing each of these five genes is shown for Cd81 (B), Cd9 (C), Cd63 (D), Sdc4 (E) and Sdcbp (F). G) To determine aggregate expression of EV genes across all cell populations, Z-scores were calculated for mean FPKM then plotted against the number of EV genes expressed @ > 50 FPKM. n= 44,779 cells with reported identity from 117 tissues of the *Tabula Muris*.

Our next goal was to compare the expression of each of these five highly abundant EV cargo genes across cell populations. Cell populations with very high expression of Cd81 (**Figure 2B**) included multiple cell types in the brain (Bergmann glial cells, oligodendrocytes, microglia, oligodendrocyte precursor cells, astrocytes and macrophages), whereas cell populations with very low Cd81 expression included multiple populations of natural killer cells and acinar cells of the pancreas. In contrast to Cd81, Cd9 was very highly expressed in multiple cell types that support innate immunity (**Figure 2C**), including multiple basal and epithelial “barrier” cell populations as well as multiple stem cell populations—although similar to Cd81, Cd9 expression was very low in acinar cells of the pancreas, as well as in multiple B cell populations. Cd63 was most abundant in stem/progenitor cell populations (**Figure 2D**), especially those from skeletal muscle (SkM), and very low in multiple immune cell populations, including T cells, B cells, natural killer cells and their precursors. Sdc4 expression was also by far the highest in skeletal muscle stem cells compared to other cell populations (**Figure 2E**), and its expression was lowest in multiple bone marrow-derived cells and leukocytes of the lung. Finally, Sdcbp expression (**Figure 2F**) was greatest in immune cells and bone marrow-derived immune cell progenitors, and lowest in acinar cells of the pancreas and brush cells of the large intestine. These data reveal that the cell populations most enriched in the most abundant EV genes are not conventional secretory cell types, lending support to the idea that circulating EVs expressing common protein cargo may originate from many cells and tissues throughout the body.

Next, we identified cell populations with high or low overall expression in our entire set of EV genes by integrating two complementary approaches — one dependent and one independent of expression in other cell populations. First, we calculated z-scores for each gene across all cell populations to represent the difference in expression of each gene in each cell population relative to other cell populations while weighting each gene equally, and to allow for a sum of z-scores for all genes as an aggregate measure of EV gene expression within a cell population. Second, as an independent measure, we also calculated the number of EV genes with mean expression over 50 FPKM in each cell population. Plotting these data together provided a two-dimensional representation of overall cell population-specific EV gene expression (**Figure 2G**). Cell populations with high combined scores included: 1) cell types of the pancreas, consistent with its role as a secretory organ, 2) antigen presenting cells, consistent with their role in secreting major histocompatibility complex (MHC), a known EV cargo [32], and 3) oligodendrocytes and Bergmann glial cells of the brain, although the explanation for this finding is less clear. Cell populations with very low aggregate expression included multiple B and T cell populations. These data provide evidence that the secretory functions of supporting cells of the brain require further examination.

### Inferring dynamic and constitutive EV-associated gene function based on expression variance

EV secretion can be constitutive or dynamic (i.e., stress-induced) [33] but the transcriptional behavior of EV genes is not well understood. High and low variance in the expression of individual genes can predict whether the encoded proteins exert dynamic or constitutive functions, respectively [34]. This means that the variance in expression of specific EV genes may provide insight into their role in EV secretion. To explore this idea, we calculated the mean-adjusted variance (coefficient of variation (CV)) for each EV-associated gene across all cell populations and plotted these data against mean transcript abundance. This analysis revealed a power-law distribution (**Figure 3A**) (note data in graph plotted on log scales). That is, genes with greater mean expression tended to have lower cell population-level variance, although this as not the case for all genes. This observation has been made in other scRNA-Seq analyses [35] and broadly suggests that most highly abundant EV genes have constitutive cellular functions. However, significant differences in expression variance were observed amongst the five abundant EV cargo genes identified in **Figure 2**. Of these genes, expression variance was greater for Cd63 (**Figure 3B**) than Sdc4 (**Figure 3C**), Cd9 (**Figure 3D**), Cd81 (**Figure 3E**) or Sdcbp (**Figure 3F**). A mixed effects one-way ANOVA revealed significantly different expression variance between genes (**Figure 3G**) which suggests that most highly abundant EV genes are constitutively expressed, while some (i.e., Cd63) are more dynamically expressed.

**Figure 3:**
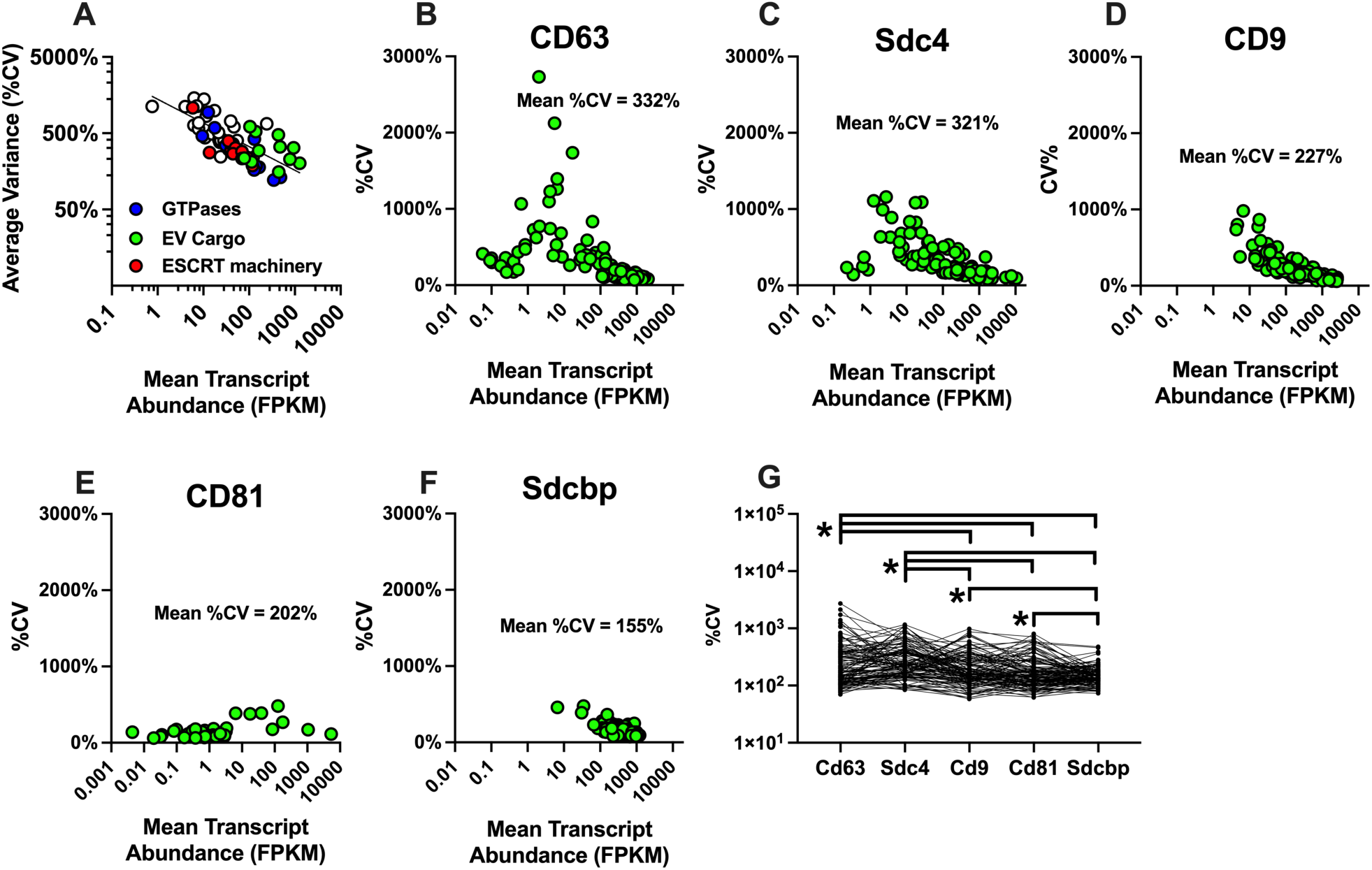
Variance of EV-associated genes within and between cell populations. A) Gene expression variance (%CV) for all EV genes plotted against mean transcript abundance. Note the data expressed in log scale. %CV of every cell population expression of CD63 (B), Sdc4 (C), Cd9 (D), Cd81 (E) and Sdcbp (F). In B-F, mean %CV across all cell populations is inset. G) Mixed-effects one-way ANOVA comparing cell population %CV. n= 44,779 cells with reported identity from 117 tissues of the *Tabula Muris*.

### Co-expression patterns of abundant EV cargo suggest differential regulatory mechanisms

EV secretion requires vesicle biogenesis, endosome processing, cargo selection and exocytosis, and each of these processes is coordinated by multiple genes. To identify putative functional interactions among EV genes and cargo, we generated gene co-expression matrices [36, 37] to determine which EV genes are most often co-expressed with abundant EV cargo and in which cell populations. To visualize co-expression data for abundant EV cargo genes, we generated heatmaps with data distributed via 2-D hierarchal clustering based on Euclidian distance and Ward agglomeration. For each EV cargo gene, we show the entire map (**Figure 4**) and clusters of highly co-expressed genes and cell populations based on the first or second level of dendrograms.

**Figure 4:**
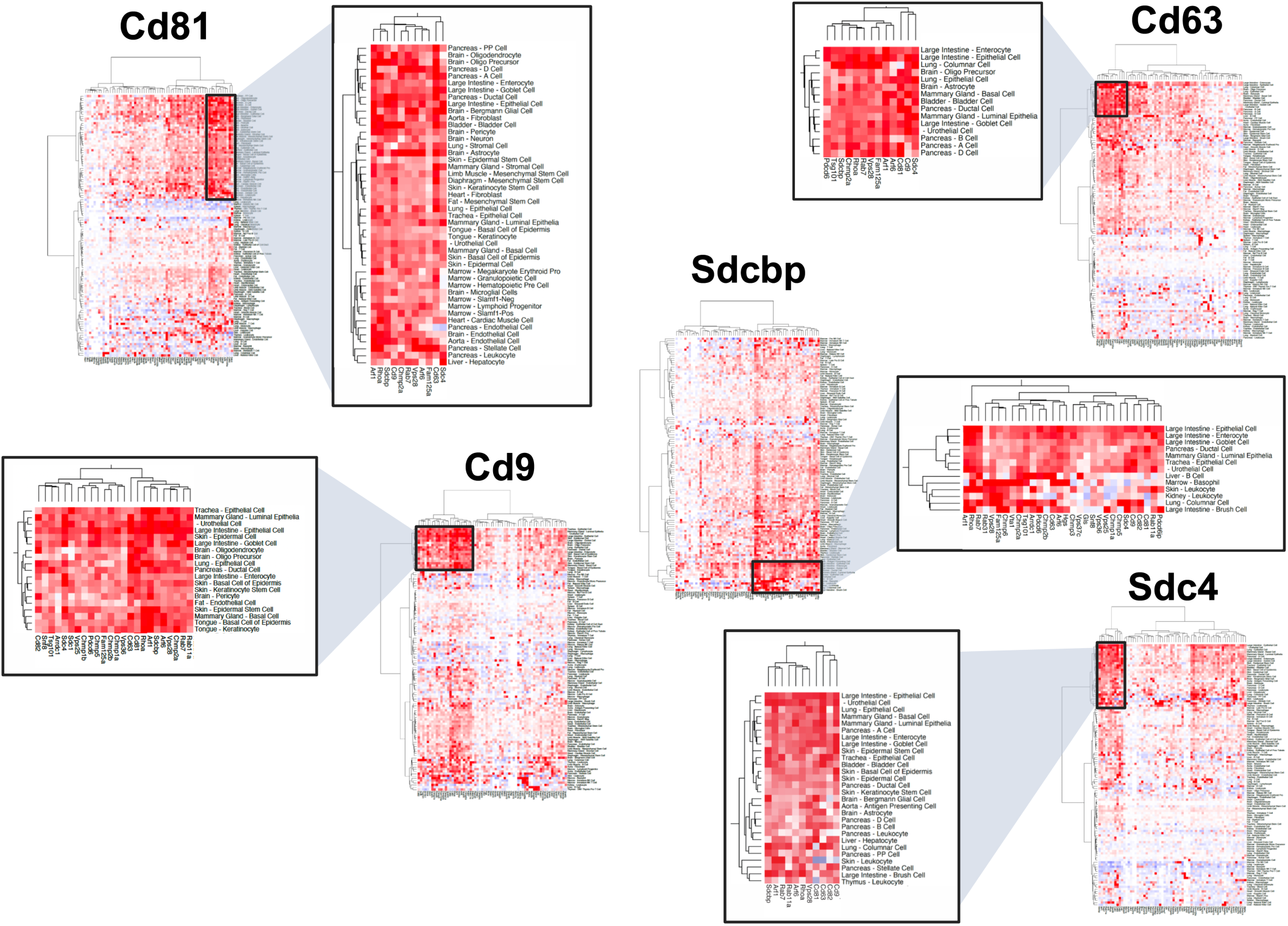
Cell population-specific co-expression of EV genes. Co-expression of EV genes encoding protein cargo (Cd81, Cd9, Cd63, Sdcbp and Sdc4) versus all other EV genes was calculated via linear regression. Heat maps with 2D hierarchical clustering were used to visual co-expression. Sub-clusters were selected from the first or second branch of each dendrogram on each axis. Heat maps generated using MD Anderson Next-Generation Clustered Heat Map (NG-CHM) tool Version 2.20.0 and figure assembled with Biorender.

We found that Cd81 was most frequently co-expressed with other abundant EV genes (i.e., Cd63, Cd9, Rhoa, Sdc4, Sdcbp and Arf1) as well as other EV genes important for endosome formation and cargo sorting (i.e., Chmp2a, Rab7, Vps28, Fam125a and Arf6). Cluster analysis revealed that the cell populations that most robustly co-expressed Cd81 with other EV genes included secretory cells of the pancreas and large intestine, barrier cells of the skin and lung, most cell types of the brain, and stem cells of the skeletal muscle and adipose. Cd9 co-expression analysis generated a larger cluster of genes than Cd81 which included every gene in the cluster identified for Cd81, along with additional genes involved in endosome formation and cargo sorting (Chmp1a, Chmp1b, Chmp2b, Chmp5, Arrdc1, Tsg101, Pdcd6, Rab11a, Snf8, Vps25 and Vps36) and EV cargo (Cd82 and Sdc4). Cell populations with robust Cd9 co-expression included cell types of the brain, and cell populations that form the physical barrier needed for innate immunity (i.e., epithelial cells and keratinocytes). Cd63 is co-expressed with many of the same genes as Cd9 and Cd81. Cell populations clustered by high Cd63 co-expression include secretory cells (similar to Cd81) and epithelial cells (similar to Cd9). Notably, however, SkM progenitor (and other stem) cells which express very high levels of Cd63 were found in a separate cluster of cells. Sdcbp co-expression clustering was more apparent by gene than by cell population. The widespread and constitutive expression of Sdcbp in many cell populations may explain this observation. Indeed, a large number of genes were frequently co-expressed with Sdcbp, including all of the co-expressed genes described above and some additional genes, including Hgs, Vta1 and Chmp6. Finally, Sdc4 was co-expressed with tetraspanins (Cd9, Cd63, Cd81 and Cd82), GTPases and Sdcbp. Cell populations that demonstrated these co-expression patterns include barrier cells (like Cd9) but not stem cells (like Cd63/81). Moreover, cell population clustering of Sdc4 co-expression also includes some cell populations (i.e., hepatocytes) that did not emerge from cell population clustering based on tetraspanin co-expression.

## DISCUSSION

It is canon in the field of EV biology to state that all eukaryotic cells can secrete EVs, but this claim is impractical, if not impossible, to directly test due to technical limitations which preclude the culture and study of some cell populations. Moreover, this claim diminishes the consistently demonstrated heterogeneity in the ability of different cells to secrete EVs and their varying cargo. Addressing this claim and the heterogeneity of EV biology between cell populations with empirical evidence meets at least two needs in the field of EV biology. First, the quantity and composition of EVs are the primary determinants of a cell population’s ability to communicate via EVs, yet no studies to date have been able to map this heterogeneity at the level of the organism. Second, predicting the ability of rare or technically challenging cell populations to secrete EVs may reveal disease etiology which could serve as a catalyst for future studies to understand these cells in greater detail. Here, our analysis of EV gene expression in the Tabula Muris predicts cell populations that secrete many (or few) EVs and reveals unique expression patterns of EV genes among cell populations.

Based on EV gene transcript abundance, our analysis supports the claim that most cell populations are capable of secreting EVs. Abundant and important EV cargo genes are enriched in different cell populations, some are dynamically expressed while others are constitutive, and co-expression analysis identifies a common set of machinery to support secretion of these cargo. Below we explore the implications of these observations in different tissues and physiological contexts.

### Observation 1: EV cargo genes are highly but differentially expressed across cell populations

Our analysis shows that the most abundant EV genes encode for the most abundant proteins found on circulating EVs. This observation is notable because it aligns with previous work showing that the expression of EV genes in cultured cells and tissues predicts the capacity of those cells and tissues to secrete EVs containing those proteins [2, 3]. These findings suggest that EV gene transcript levels may predict EV protein cargo secretion.

For example, robust gene expression of Cd9 and Cd81 combined with their abundance in circulating EVs together suggest that constitutive exchange of Cd9^+^ and Cd81^+^ EVs between cells may occur *in vivo.* Our findings here further support this idea by illustrating the heterogeneity of Cd81 (**Figure 1D**) and Cd9 (**Figure 1E**) expression among cell populations. This is noteworthy because multiple reports describe functional redundancy between Cd81 and Cd9 [38–40] but recent studies suggest opposing functions. For example, Cd81 preserves metabolic health in adipocytes and counteracts the effects of overnutrition by high fat feeding via anti-inflammatory signaling [41]. By contrast, Cd9 marks a population of pro-inflammatory cells in the brain [42] and adipose tissue [43]. Exercise can increase the circulating abundance of EVs expressing both Cd9^+^ and Cd81^+^ [44, 45], often on the same EV, which could be reconciled by the observation that exercise promotes both inflammation and metabolic health. The functional differences between these proteins when delivered to recipient cells via EVs are not known but could be defined using gain/loss of function experiments in cells and model organisms with naturally high or low expression of each gene.

It is also interesting that Cd81 is more abundant than Cd9 in mice across all cell populations because our group recently reported that Cd9 is more abundant than Cd81 in circulating plasma EVs in mice [2]. It is worth noting that others have made the opposite observation (Cd81 > Cd9) in human plasma [44], so circulating tetraspanin protein abundance could be species-specific. Our analysis here did not identify any blood-borne cell populations that would explain a greater abundance of circulating Cd9^+^ EVs in mice, so this suggests that there may be differences in half-life for different EV populations in biofluids or that tissue-resident cells may be major contributors to circulating EV populations.

We find that expression variance is not consistent across all abundant EV genes. Instead, we find that Cd9, Cd81 and Sdc4 are expressed more constitutively whereas Cd63 and Sdcbp are more dynamically expressed. This is notable because (before being secreted in/on EVs) Cd9 [46], Cd81 [47] and Sdc4 [48] are found almost exclusively on the surface of cells, while Cd63 [49] and Sdcbp [50] are found within the cell. Because Cd63 and Sdcbp both directly regulate EV secretion [51–54], this suggests that Cd63 and Sdcbp expression may be useful predictors of cellular EV secretion.

### Observation 2: SkM stem cells highly express genes that encode abundant EV cargo proteins

Regenerative stem cell therapies have emerged as a promising area of research due in part to the therapeutic properties of EVs secreted by stem cells [55]. Our analysis shows that SkM stem cells are enriched in EV genes like Cd63 and Sdc4, which supports the notion that these cells are a major source of EVs in the body on a per cell basis. This idea is also supported by multiple experiments by independent groups. For example, Garcia-Martin et al. performed a rigorous and direct comparison of cellular transcriptomics and EV proteomics in immortalized myocytes, adipocytes, hepatocytes and endothelial cells [3]. They demonstrated that SkM-like C2C12 myotubes have greater gene expression of tetraspanins than adipocyte-like 3T3-L1 cells and secrete far greater quantities of tetraspanin proteins in EVs [3]. Murach et al. have published a series of papers demonstrating that SkM stem cells *in vivo* secrete EVs that travel to and act upon other cells within SkM tissue to remodel the extracellular matrix and promote hypertrophy independent of stem cell fusion with existing myofibers [56, 57]. The functions of SkM stem cell EVs have been attributed to syndecans like Sdc4 [58] and microRNAs (miR 206) [57, 59], but the latter may be scarce in EVs (i.e., < 1 miRNA / EV) [60]. This suggests that the large quantity of EVs secreted by SkM stem cells and/or preferential targeting to specific recipient cell populations could enable these EVs to exert clinically significant effects. The secretion of EVs by SkM stem cells is also consistent with the large quantities of EVs secreted by mature SkM myofibers [2] (which were not collected in the *Tabula Muris*) and the systemic benefits of exercise [61].

An acute bout of exercise increases the circulating abundance of tetraspanin-expressing EVs [44, 45] and while SkM is a logical source for some of these EVs, their cellular origin(s) have not been well defined experimentally. What is known is that SkM myofibers secrete far more EVs than the same mass of adipose tissue [2], these EVs can be found within SkM tissues [62] as well as the circulation [2] and they accumulate in distant recipient tissues like white adipose during exercise where they enhance lipolysis [63]. Since SkM tissue is also the largest tissue (by mass) in non-obese humans, this evidence places SkM as one of the most abundant sources of EVs in the entire body. Future studies could build upon this existing knowledge by quantifying the circulating abundance and tissue accumulation of protein-specific SkM EV populations *in vivo* using commercially available Cd9 [64] and Cd63 [65] EV reporter mice and/or novel Cd81 EV reporter mice [66]. If SkM is truly a major source of circulating EVs, then the manipulation of SkM EVs could be a pharmacological strategy to improve human health (i.e., mimicking exercise) and an ideal model to study mechanisms governing interorgan EV trafficking and recipient cell function.

### Observation 3: Cd81 is a primary EV cargo trafficked within the brain

EV biology of the brain has garnered attention in recent years in part due to the ability of EVs to traffic pathological proteins like tau and amyloid beta between cells [64, 67–70]. For this reason, EV secretion from cells of the brain may influence the development and progression of neurodegenerative diseases like Alzheimer’s. We found that multiple cell populations of the brain have high expression of Cd81 but low expression of Cd9. This finding suggests that large quantities of Cd81^+^ EVs should be present in brain tissue relative to Cd9^+^ EVs. Indeed, a recent study demonstrated that EVs obtained from the human brain interstitium express more Cd81 than Cd9 [71]. It is not yet clear to what extent EVs are traveling to and from different cell populations within the brain, so the horizontal transfer of pathogenic proteins remains undefined. However, what is clear from our analysis is that Cd81^+^ EVs in the brain could originate from multiple cell populations. Despite the high abundance of Cd81 mRNA in cells of the brain and Cd81^+^ EVs in the brain interstitium, very little is known regarding the function of Cd81 in the brain and nothing is known about the function of Cd81^+^ EVs trafficked within or beyond the brain. Whole body Cd81 knockout mice are characterized by increased number of glia [72] and there is some evidence that Cd81^+^ neuron-derived EVs are reduced in Alzheimer’s disease [73, 74], but no cell type-specific effects of Cd81 have been examined or described. Functional Cd81 protein can be transferred between cells [75], so the directional trafficking of CD81^+^ EVs within the brain could be explored using different variations (e.g., Cre drivers) of a novel Cd81 EV reporter mouse model [66] to elucidate disease modifying effects during cognitive decline.

### Observation 4: EV gene expression in rare cell populations suggests novel modes of communication

While there are a multitude of abundant cell populations that are relatively easy to culture and study *in vitro*, there are also cell populations that are not amenable to the cell culture conditions required to collect and analyze secreted EVs. In these rare and difficult cell populations, analysis of EV gene expression may be one of the only feasible strategies to predict how these cells might contribute to EV-based communication. To demonstrate the possible utility of this approach we highlight Bergmann glial cells of the brain. Bergmann glia reside in the cerebellum where they support cerebellar morphogenesis and homeostasis [76] and play a particularly important role in the support of Purkinje neurons. Here we show that Bergmann glia demonstrate the greatest mean expression of Cd81 of any cell population, and have high aggregate expression of EV genes, yet their role in EV biology has not been previously described. Interestingly, Bergmann glia express Cd9 at low levels, which further suggests that Cd81 is likely a primary EV cargo secreted by these cells. Moreover, some data suggest an underappreciated role of Purkinje neuron/cerebellar degeneration early in Alzheimer’s disease progression [77], suggesting that the EV-related functions of these cells warrant further study. Because EV gene expression predicts EV secretion and cargo in easily cultured immortalized cell lines [3], it is possible that single-cell transcriptomics will be a useful approach to understanding EV biology in rare cell populations like these.

### Limitations

There are at least two limitations to this project. First, the majority of genes analyzed have functions that may be unrelated to EV biology. So, although the expression of EV genes is associated with EV secretion in cultured cells and tissues, it is unclear whether these EV genes play comparable roles in each cell population and whether the proteins encoded by EV cargo genes are actually trafficked into EVs at a similar rate/density/etc. in different cell populations. Second, gene expression does not always align with protein expression, and this is potentially complicated by the nature of single-cell RNA-seq data, in which some genes/transcripts are detected more robustly than others. That is, the extent to which transcript abundance is an accurate reflection of the cellular protein machinery required to form and secrete EVs is unclear but our findings here enable the generation of hypotheses to answer this question. Moreover, advances in single cell proteomics should facilitate those studies in the near future.

### Conclusions

Our analysis offers new, comprehensive insight into the differential transcriptional expression of genes encoding the machinery and cargo of EVs. We have identified cell populations that are predicted to secrete large quantities of EVs and, in some cases, large quantities of specific EV cargo. We explore the physiological implications and existing literature in which EVs were studied directly and emphasize the need to experimentally validate the hypotheses generated from our analysis. We are hopeful that this resource can serve as a starting point for investigators with an interest in EV biology to strengthen the premise of their studies by establishing the extent to which their cell populations of interest may secrete EVs and, in this way, advance our understanding of endocrinology and organ systems physiology.

## METHODS

### Source Data

FACS-sorted single cell mRNA transcriptome data (reported in FPKM) from the *Tabula Muris* [5] was obtained via Figshare (https://figshare.com/projects/Tabula_Muris_Transcriptomic_characterization_of_20_organs_and_tissues_from_Mus_musculus_at_single_cell_resolution/27733). Only cells with a defined origin in the original report were studied in this project (n= 44,779 cells). A total of 117 distinct cell populations (cell type x tissue) were identified and analyzed. A notable strength of the *Tabula Muris* dataset is that the investigators used fluorescent antibodies to enrich for rare cell populations. Detailed methods regarding sample collection, FACS sorting and single cell RNA-Seq workflow (including cell identification) can be found in their original publication.

### Selection of EV Genes

67 EV-associated genes were selected based on recent expert reviews[7] and/or structural similarity to experimentally validated genes/proteins. A detailed explanation of the genes selected is provided in **Results**.

### Z-score adjustment

To examine cell population-specific expression of EV-associated genes, we adjusted the expression of each gene based on Z-score:

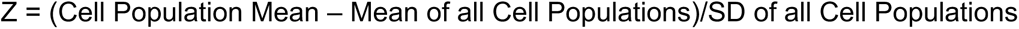

Z score adjustment normalizes expression of each gene relative to all cell populations. This adjustment allowed us to compare the amplitude of expression differences across all EV genes and cell populations. Although expression of some EV-associated genes are not normally distributed, Z-score normalization is still appropriate in this case because the central limit theorem states that normality is not necessary with a sufficiently large sample size (i.e., N > 30) [78].

### Gene Co-Expression Analyses

We calculated single cell gene co-expression between our five highly abundant EV cargo genes (Cd9, Cd63, Cd81, Sdc4 and Sdcbp) with every other EV gene (n=67) within every cell population (n=117) via linear regression using Graphpad Prism. We visualized gene co-expression patterns via hierarchical clustering based on Euclidean distance and Ward agglomeration using the MD Anderson Next Generation Clustered Heat Map (NG-CHM) Builder Version 2.20.0 (https://www.ngchm.net/) [79, 80]. The heat map color scale was standardized from 0.7 (solid red) to 0 (white) to -0.7 (solid blue). Clusters were emphasized based on strong co-expression at the first or second level of each dendrogram on each axis.

## ACKNOWLEDGEMENTS

We would like to acknowledge the Tabula Muris Consortium and the Chan-Zuckerberg Initiative for the generation and sharing of the *Tabula Muris* dataset.

## Competing Interests

No competing interests are declared by the authors.

## Funding Information

American Heart Association (IPA1834110052) to D.S.L. and National Institutes of Health (AG078859) to T.J.L.

## Notes

### Competing Interest Statement

The authors have declared no competing interest.

